# Dietary Branched Chain Amino Acids Modify Post-Infarct Cardiac Remodeling and Function in the Murine Heart

**DOI:** 10.1101/2024.07.12.603348

**Authors:** Daniel C. Nguyen, Collin K. Wells, Madison S. Taylor, Yania Martinez-Ondaro, Kenneth R. Brittian, Robert E. Brainard, Joseph B. Moore, Bradford G. Hill

**Author notes:** **Correspondence: Bradford G. Hill**, **Ph.D.**, Center for Cardiometabolic Science, Christina Lee Brown Envirome Institute, Division of Environmental Medicine, Department of Medicine, University of Louisville, 580 S. Preston St., Rm 321E, Louisville, KY 40202.

## Abstract

**Introduction:** Branch-chain amino acids (BCAA) are markedly elevated in the heart following myocardial infarction (MI) in both humans and animal models. Nevertheless, it remains unclear whether dietary BCAA levels influence post-MI remodeling. We hypothesize that lowering dietary BCAA levels prevents adverse cardiac remodeling after MI.

**Methods and Results:** To assess whether altering dietary BCAA levels would impact circulating BCAA concentrations, mice were fed a low (1/3×), normal (1×), or high (2×) BCAA diet over a 7-day period. We found that mice fed the low BCAA diet had >2-fold lower circulating BCAA concentrations when compared with normal and high BCAA diet feeding strategies; notably, the high BCAA diet did not further increase BCAA levels over the normal chow diet. To investigate the impact of dietary BCAAs on cardiac remodeling and function after MI, male and female mice were fed either the low or high BCAA diet for 2 wk prior to MI and for 4 wk after MI. Although body weights or heart masses were not different in female mice fed the custom diets, male mice fed the high BCAA diet had significantly higher body and heart masses than those on the low BCAA diet. Echocardiographic assessments revealed that the low BCAA diet preserved stroke volume and cardiac output for the duration of the study, while the high BCAA diet led to progressive decreases in cardiac function. Although no discernible differences in cardiac fibrosis, scar collagen topography, or cardiomyocyte cross-sectional area were found between the dietary groups, male mice fed the high BCAA diet showed longer cardiomyocytes and higher capillary density compared with the low BCAA group.

**Conclusions:** Provision of a diet low in BCAAs to mice mitigates eccentric cardiomyocyte remodeling and loss of cardiac function after MI, with dietary effects more prominent in males.

## INTRODUCTION

The cardiac response to injury involves coordination of several responses to repair damaged tissue. Particularly important are metabolic changes that influence the heart’s ability to maintain sufficient energy to fulfill work demands and activation of responses that facilitate limited cardiac repair. Several metabolic pathways and metabolites are implicated in post-infarction remodeling, with metabolic pathways such as glycolysis, fatty acid oxidation, and ketone body oxidation contributing to changes in the phenotype and function of both parenchymal and mesenchymal cells of the heart.^1–5^ Interestingly, recent studies have shown that branched chain amino acids (BCAAs) are not only linked with cardiovascular disease risk in humans (reviewed in ^6^), but are also elevated substantially in the context of heart failure in both clinical^7^ and preclinical^8, 9^ scenarios. Nevertheless, it remains unclear whether elevations in BCAAs are deleterious to cardiac healing and function after infarction and whether modulating BCAA levels could be an actionable strategy for improving outcomes.

BCAAs, as essential amino acids, are primarily obtained from the diet and play crucial roles in protein synthesis and energy homeostasis. Although BCAAs are thought to be required for anabolic responses that contribute to repair of tissue in response to physiological stress (e.g., exercise),^10^ chronically high levels of circulating BCAAs are associated with insulin resistance and cardiometabolic disease^11^ and could act as mediators of heart failure pathogenesis.^6, 12^ Indeed, modifying dietary BCAA intake has been shown to regulate metabolic health, leading to improvements in glucose tolerance and body composition.^13^ However, their role in cardiac remodeling after myocardial infarction (MI) remains unclear. Some studies have reported that consumption of an essential amino acid-enriched diet and a high-leucine diet can attenuate pathological adaptation in different models of cardiac injury.^14, 15^ Conversely, other studies have found that, in the context of pressure overload, a BCAA-enriched meal can accelerate cardiac dysfunction, whereas a BCAA-free diet may mitigate maladaptive hypertrophy and fibrosis.^16, 17^ Notably, few efforts have been made to interrogate directly whether dietary BCAAs influence remodeling following MI.

To address this issue, we examined in mice whether dietary alterations in BCAA levels could influence remodeling and function after MI. We found that provision of a diet low in BCAAs is sufficient to improve cardiac function in both male and female mice after MI. Low dietary BCAAs led to sustained cardiac output after MI, whereas high dietary BCAAs promoted progressive loss of cardiac function, associated with eccentric cardiomyocyte hypertrophy. Notably, the effect of dietary BCAAs on remodeling was more pronounced in male mice. These findings indicate that dietary changes in BCAA composition influence cardiac structure and function after MI, lending support to the idea that careful modulation of the diet after MI could be an actionable strategy to optimize cardiac recovery.

## MATERIALS AND METHODS

### Experimental animals and diets

Experimental protocols were reviewed and approved by the University of Louisville Institutional Animal Care and Use Committee, and all animal studies were completed in compliance with the *Guide for the Care and Use of Laboratory Animals*. Male and female C57BL/6J mice were purchased from the Jackson Laboratory. The low (Cat: TD.150662) and high (Cat: TD.170323) BCAA custom modified diets were purchased from Envigo Teklad (see supplementary files for diet formulations). The standard chow diet was purchased from LabDiet (Cat: 5010, see supplementary files for diet formulations). All animals were housed in a pathogen-free facility under controlled conditions (24°C, 44–65% relative humidity, 12-hour light/dark cycle).

### Rigor, transparency, and reproducibility

The study was conducted in accordance with our recently published guidelines on scientific rigor.^18^ Based on prior experiences and subsequent power analyses, a final “n” of 9 was the anticipated goal for this study. As such, 11 female and 13 male mice were used for each diet group with the expectation of post-MI attrition being different between sexes. Echocardiographic confirmation of infarction was required for inclusion in subsequent analysis. To maintain a rigorous experimental design, *in vivo* experiments included blinding of surgeons, sonographers, and microscopists. Tissue sample processing and analysis were performed in a blinded manner as well.

### Non-reperfused myocardial infarction and echocardiography

Adult, 12-week-old, male and female mice were subjected to non-reperfused myocardial infarction (MI), as described.^19, 20^ Briefly, mice were first anesthetized through intra-peritoneal injections of (pentobarbital, 50 mg/kg; ketamine, 50 mg/kg) before surgery. Subsequently, they were orally intubated and ventilated with oxygen. Using a 7-0 silk suture, the left coronary artery was permanently ligated and then the chest wall was sutured closed. Mice were extubated only after the recovery of spontaneous breathing, and analgesia (meloxicam, 10 mg/kg) was provided immediately prior to surgery as well as twice a day at 24 and 48 h post-surgery. Cardiac structure and function were assessed through transthoracic echocardiography (Vevo 3100, Visual Sonics) at the indicated times.

### Histopathology

At the conclusion of the study, hearts were excised and arrested in diastole with ice-cold 2% potassium chloride. Each heart was then sectioned into 1-mm cross-sectional segments and fixed in 10% formalin prior to being embedded in paraffin, sectioned (4 µm), and mounted. Slides were then deparaffinized and rehydrated before staining. Picrosirius red staining (Picric Acid, Sigma, P6744; Direct Red 80, Sigma, 365548) was used to detect collagen in intact sections. Cardiomyocyte size was assessed by staining with wheat germ agglutinin (WGA, Vector Labs, RL-1022) to demarcate cell membranes followed by assessment of cardiomyocyte cross-sectional area^21^ and length.^22, 23^ Capillary density was measured by staining with isolectin B4 (Vector Labs, FL-1201). DAPI (Invitrogen, D3571) was used to detect nuclei. Sections were visualized using the Keyence Imaging System. Acquired images were then analyzed using the Keyence BZ-X800 analyzer.

### Second harmonic generation (SHG) imaging

Transverse myocardial tissue sections were deparaffinized by heating at 80°C for 30 min, followed by rinsing in xylene. The sections were then rehydrated stepwise in decreasing concentrations of ethanol (100%, 96%, 90%, 80%), with a final submersion in deionized water for 3 min. After a brief placement in 1× PBS, the sections underwent second harmonic generation (SHG) imaging using a Nikon A1R MP+ multiphoton microscope with an Apo LWD 25× immersion objective. The excitation laser was tuned to 920 nm, and the SHG signal was collected through a DAPI bandpass filter with an emission wavelength of 446 nm. Images were acquired with a resolution of 3595 × 3598 pixels at a scan speed of 15 s. Over 20 regions of interest (ROI: 256 × 256 pixels) were selected per heart, focusing on the longitudinal aspect of collagen fibers within infarct scar regions. The ROIs were subject to computational analyses using MATLAB software packages CurveAlign and CT-FIRE,^24^ as described,^25^ to quantify macrostructural attributes of the collagen fiber network (collagen fiber alignment, width, and straightness).

### Measurement of plasma BCAAs

Blood samples were collected via a right ventricular puncture using a 23-gauge needle and an EDTA-coated syringe. Samples were centrifuged at 3,500*g* for 20 min, followed by separation of the plasma fraction. Plasma BCAA levels were then measured using a commercially available BCAA assay kit (Abcam; ab83374), according to manufacturer’s protocol.

### Primary Cardiac Fibroblast Isolation and Culture

Cardiac fibroblasts were isolated from naïve, adult mice. Briefly, following euthanasia, hearts were excised, and the aorta was cannulated with a 23-gauge needle. Ice-cold PBS was flushed through the aorta, and excess aorta and adipose tissue were trimmed from the heart prior to mincing with a razor blade under sterile conditions. Heart tissue was then digested in collagenase solution for 45 min at 37°C with gentle agitation. The collagenase solution contained type II collagenase (Worthington, 46H16739) prepared in sterile PBS at 5000 units per 7.5 ml per heart. Subsequently, culture medium [i.e., Dulbecco’s Modified Eagle’s Medium/F12 GlutaMax (DMEM, Gibco 10565018) containing 10% fetal bovine serum (FBS, VWR 1500-500H), 1% penicillin/streptomycin/amphotericin B (Sigma, A5955), 1% insulin transferrin selenium (ITS-G, Gibco 41400045), and 20 ng/ml human bFGF (Peprotech 100-18B)] was added to quench collagenase activity, and the mixture was centrifuged at 500*g* for 10 min. Supernatants were aspirated, and cell pellets were resuspended in culture media before being passed through a 70 µm cell strainer. Cells were then plated in a T-75 cell culture flask and incubated at 37°C in 5% CO_2_. Media was changed after 2 h and again after 24 h.

Cardiac fibroblasts were used at passage 1 and serum-starved in custom BCAA-free DMEM/F12 GlutaMax media (ThermoFisher) without any supplementation for 24 h before treatment with TGFβ (10 ng/ml, PeproTech, 100-21) for 48 h. To modify individual BCAA concentrations, L-leucine (Sigma-Aldrich, L8912), L-isoleucine (Sigma-Aldrich, I7403), and L-valine (Sigma-Aldrich, V0513) were added individually, as indicated. The concentrations used (100μM, 200 µM, 400μM) included 45.9% valine, 35.3% leucine, and 18.8% isoleucine, similar to the ratio occurring in the murine heart.^8^

### Western Blot Analysis

Total protein was harvested after washing cells with ice-cold PBS. Cells were scraped with a rubber policeman in lysis buffer (20 mM HEPES, 110 mM KCl, 1 mM EDTA, 1% IGE-PAL, and 0.1% SDS, pH 7.0) containing protease inhibitor cocktail (Sigma-Aldrich, P8340). The cell lysates were incubated on ice for 30 min and then centrifuged at 20,000*g* for 20 min. The supernatants were then carefully removed and saved for total protein measurements (Lowry assay) and Western blotting.

Protein samples were mixed with Laemmli sample buffer (125 mM Tris-HCl, 10% SDS, 50% Glycerol, 0.05% Bromophenol Blue, pH 6.8) and incubated at 95°C for 5 min. Each sample was then loaded on a 7.5% acrylamide/bis SDS-PAGE gel for electrophoresis at 120 V. SDS-PAGE-resolved proteins were then transferred overnight onto a PVDF membrane (Cytiva, 10600021) at 4°C at 20 V. Subsequently, membranes were blocked for 1 h at room temperature with Tris-buffered saline containing Tween-20 (TBS-T) and 5% bovine serum albumin. Membranes were probed with primary antibodies overnight at 4°C. The following day, membranes were washed three times using TBS-T and then incubated with secondary antibody for 1.5 h. Membranes were then washed three times with TBS-T before exposure to Pierce™ ECL Plus Western Blotting Substrate (Thermo Scientific, 32132) and imaging on a ChemiDoc imager (BioRad). Protein was normalized to total lane protein with an amido black stain, and parallel blots were normalized to an anchor protein. The primary antibodies used included: anti-COL1A1 (1:20,000; Thermo Fisher, PA5-29569), anti-Periostin (1:5,000; Millipore, ABT253), and anti-α-smooth muscle actin (1:30,000; Cell Signaling, 19245S). The secondary antibody used was HRP-linked anti-rabbit IgG (1:2,500; Cell Signaling, 7074).

### Statistical Analysis

Data are expressed as mean ± SEM. Normality was assessed via Shapiro-Wilk test. Mann–Whitney U test, Unpaired 2-tailed t test, Kruskal-Wallis test with Dunn’s post-hoc, one-way ANOVA with Tukey’s post-hoc, or two-way repeated measurement ANOVA with Šidák post-hoc were performed for between group comparisons, as appropriate and as indicated in the Figure Legends. Comparisons were considered significant when P < 0.05.

## RESULTS

### A diet low in BCAAs decreases circulating BCAA concentrations in mice

To investigate whether altering dietary BCAA levels is sufficient to influence blood levels of BCAAs, we fed wild-type, male and female, C57BL/6J mice a low or high BCAA custom diet over a 7-d period (**Fig. 1A**); these diets have 1/3× or 2× the level of BCAAs, respectively, than the typical “normal chow” diet, which was included as a comparator (see **Suppl. Tables 1–3** for diet compositions). Plasma BCAA concentrations measured at the conclusion of the pilot study revealed that the high BCAA diet did not elevate BCAAs beyond that of the normal chow diet; however, the low BCAA diet decreased circulating BCAAs by more than 2-fold (**Fig. 1B**). Although the female cohort did not exhibit any differences in body mass or heart weight normalized to tibial length (HW/TL) between groups, the male cohort provided the low BCAA diet showed a trend toward lower body weight and cardiac mass (**Fig. 1C, 1D**). These results demonstrate that provision of a diet low in BCAAs effectively reduces circulating BCAA levels in mice.

**Figure 1:**
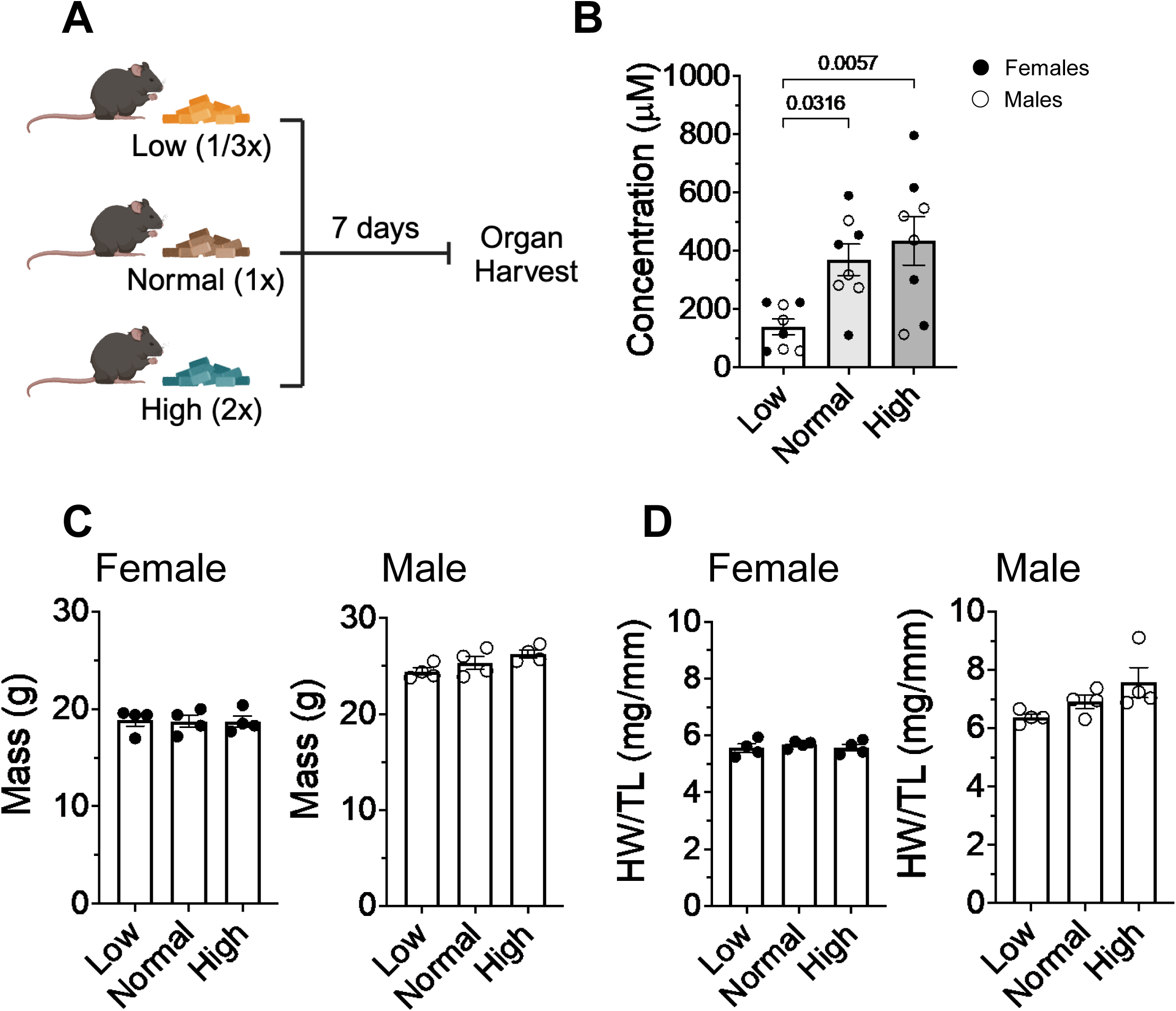
Dietary BCAA levels modulate circulating BCAAs. **A)** Mice were fed a low (1/3×), normal (1×), or high (2×) BCAA diet for 7 d prior to measurement of: (**B**) plasma BCAA concentrations; (**C**) body mass; and (**D**) heart weight (normalized to tibial length; HW/TL). n = 4 male and 4 female mice per group. Panel B shows pooled male and female mice. Data are mean ± SEM. Normality was assessed via Shapiro-Wilk test, Panel B–D, one-way ANOVA with Tukey’s post-hoc or Kruskal-Wallis with Dunn’s post-hoc. HW = heart weight, TL = tibia length

### BCAAs influence body mass and cardiac size in a sex-dependent manner

To assess the chronic effects of modified BCAA diets in a cardiac injury model, male and female mice were fed either a low BCAA diet or a high BCAA diet for 2 weeks prior to, and for 4 weeks after, MI (**Fig. 2A**). We did not include a normal chow dietary group because the composition of the normal chow is vastly different from the BCAA custom diets, which were otherwise matched for nitrogen, carbohydrate, fat, and calorific content (**Suppl. Tables 1–3**), and because the normal and high BCAA diets led to similar circulating BCAA concentrations (Fig. 1B). Baseline body weights were measured 1 week prior to surgery, and at 2- and 4-weeks post-MI. Consistent with our previous findings in the acute feeding pilot study, the female cohort did not display differences in body weight (**Fig. 2B**). Conversely, male mice fed a high BCAA diet consistently demonstrated higher body weights compared with mice fed a low BCAA diet (**Fig. 2C**). Additionally, gravimetric assessments 4 weeks post MI revealed that male mice fed the low BCAA-fed had significantly lower heart mass (HW/TL) and wet/dry lung weight ratios (**Fig. 2D, 2E**). Notably, these effects were not observed among the female cohort, suggesting that dietary BCAA levels significantly influence body weight, heart mass, and wet/dry lung weight ratio in a sex-dependent manner.

**Figure 2:**
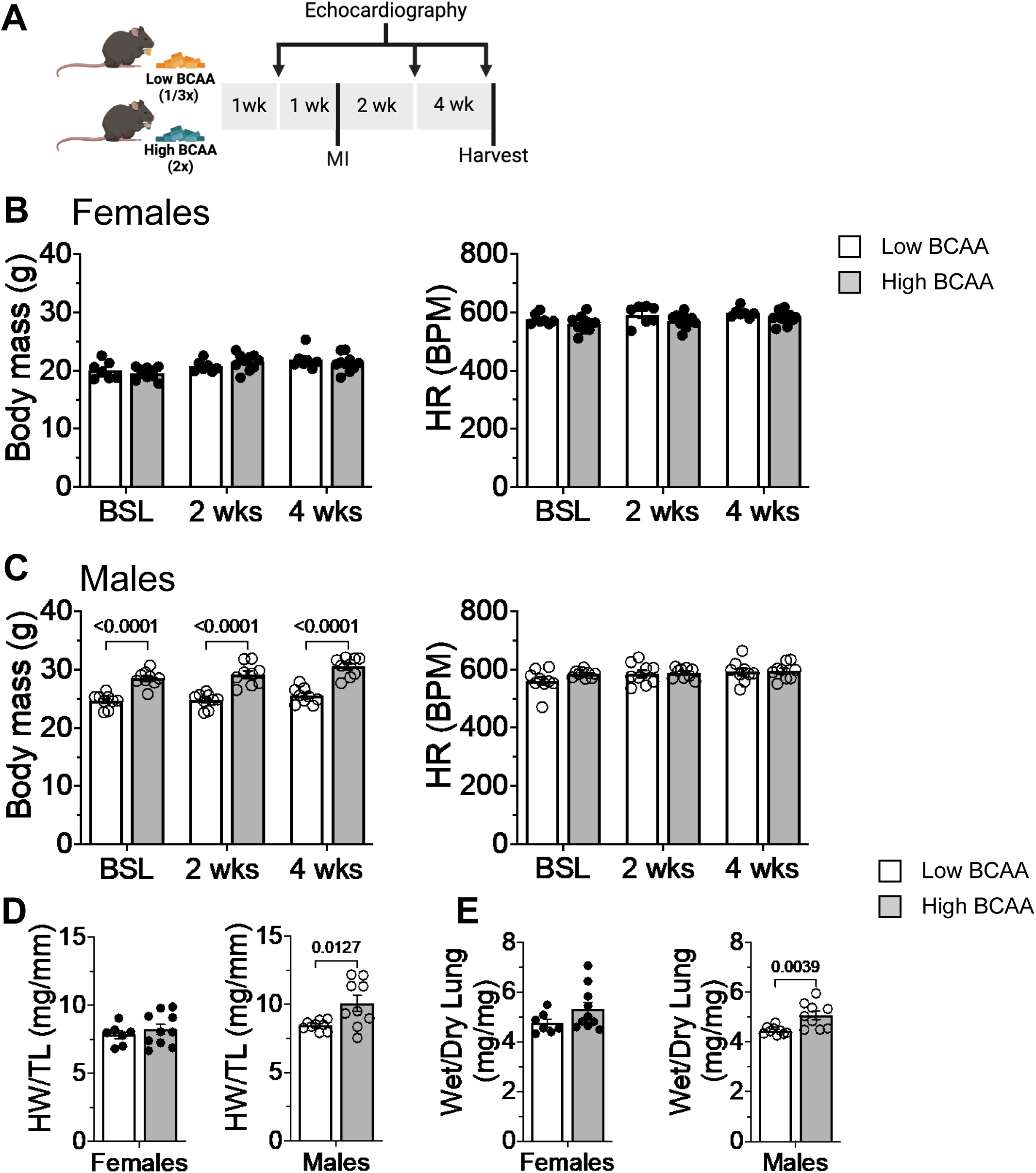
Low dietary BCAA levels prevent cardiac hypertrophy and MI-induced congestive heart failure. Gravimetric measurements: (**A**) Schematic of study design: mice were fed a low or high BCAA diet for 7 d prior to non-reperfused MI and continued on the respective diet for 28 d after MI; (**B**) Female and (**C**) male body mass and heart rate measurements recorded at baseline (BSL), 2 wk, and 4 wk post-MI; (**D**) Heart weight normalized to tibia length (HL/TL) measured 4 wk post-MI; (**E**) Wet/dry lung mass ratio measured 4 wk post-MI; n = 7–10 mice per group. Data are mean ± SEM. Normality was assessed via Shapiro-Wilk test Panel B-C, two-way repeated measures ANOVA with Šidák post-hoc; Panel D-E, Unpaired 2-tailed t-test; HW = heart weight, TL = tibia length.

### Lower dietary BCAA consumption prevents worsening of cardiac function after infarction

To study the effects of dietary BCAAs on MI-induced cardiac dysfunction and remodeling, mice were subjected to echocardiographic assessment. Serial echocardiograms were collected immediately prior to MI (baseline) and at 2- and 4-weeks post-MI (see Fig. 2A). We first examined indices of cardiac function with males and females grouped together. Although we found that dietary BCAAs did not significantly influence several volumetric and functional parameters (e.g., ventricular volumes (LVEDV and LVESV), ejection fraction (EF), fractional shortening (FS), isovolumetric relaxation time (IVRT); **Fig. 3A– 3E**), we found that they had more prominent effects on parameters that are calculated from aggregate volumetric indices, i.e., stroke volume (SV), cardiac output (CO), and cardiac index (CI). After MI, mice fed the high BCAA diet showed progressive deterioration in function, marked by reductions in SV, CO, and CI. In contrast, the low BCAA diet preserved these measures of ventricular performance (**Fig. 3F–3H**). Furthermore, mice fed the high BCAA diet had larger left ventricles (LV), as assessed by calculated LV mass (**Fig. 4A**), with no appreciable differences observed in LV internal diameters (**Fig. 4B**). Consistent with their larger LV mass, mice fed the high BCAA diet had thicker anterior and posterior wall measurements during systole and diastole (**Fig. 4C, 4D**). These findings indicate that a diet low in BCAAs prevents LV hypertrophy after MI in mice.

**Figure 3:**
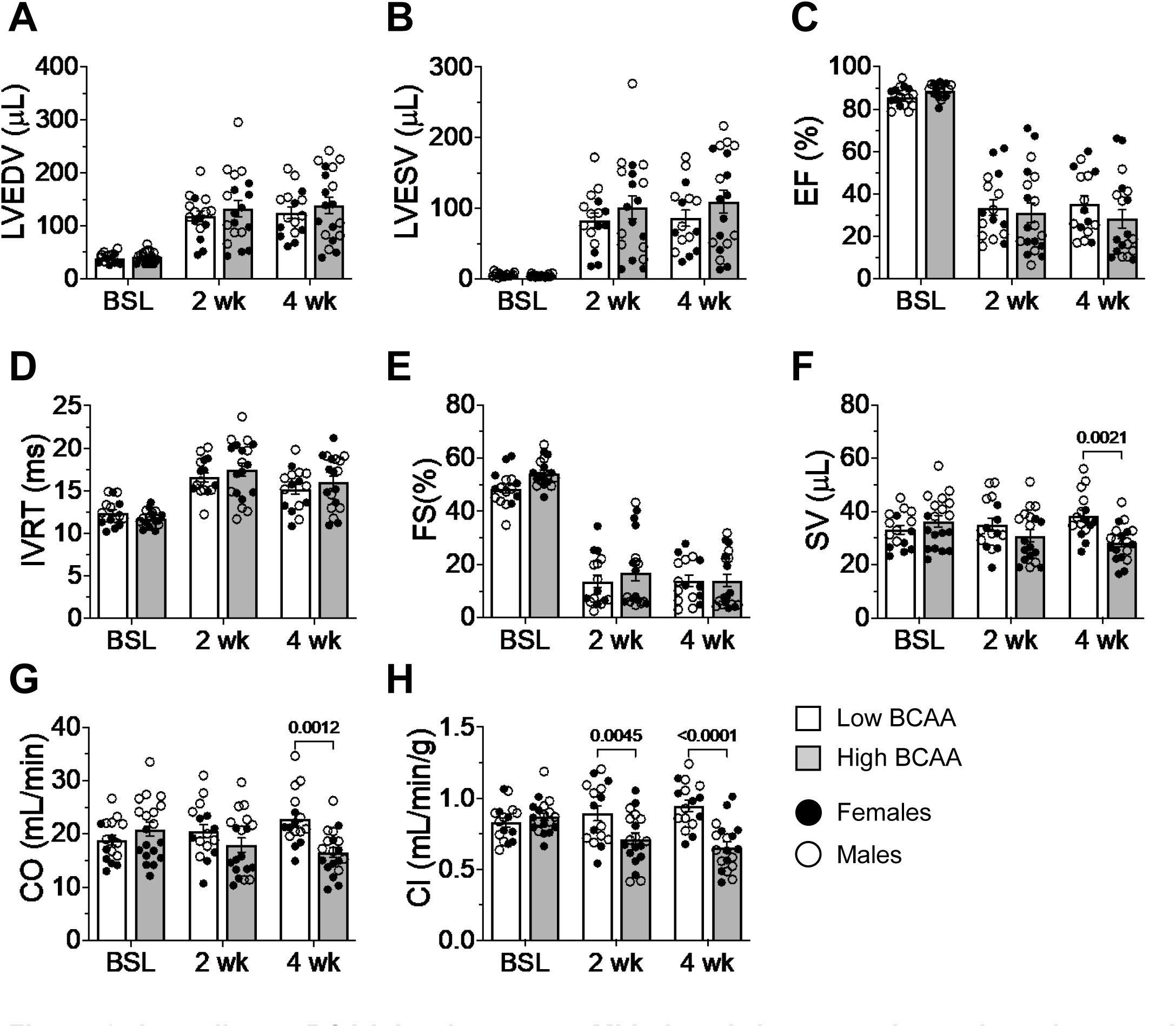
Low dietary BCAA levels prevent MI-induced decreases in stroke volume and cardiac output. Echocardiographic measurements for: (**A**) End diastolic volume (EDV); (**B**) End systolic volume (ESV); (**C**) Ejection fraction (EF) (**D**) Isovolumetric relaxation time (IVRT); (**E**) Stroke volume (SV); (**F**) Fractional shortening (FS); (**G**) Cardiac output (CO); (**H**) Cardiac index (CI); n = 16–19/group, males and females pooled. Data are mean ± SEM. Two-way repeated measures ANOVA with Šidák post-hoc.

**Figure 4:**
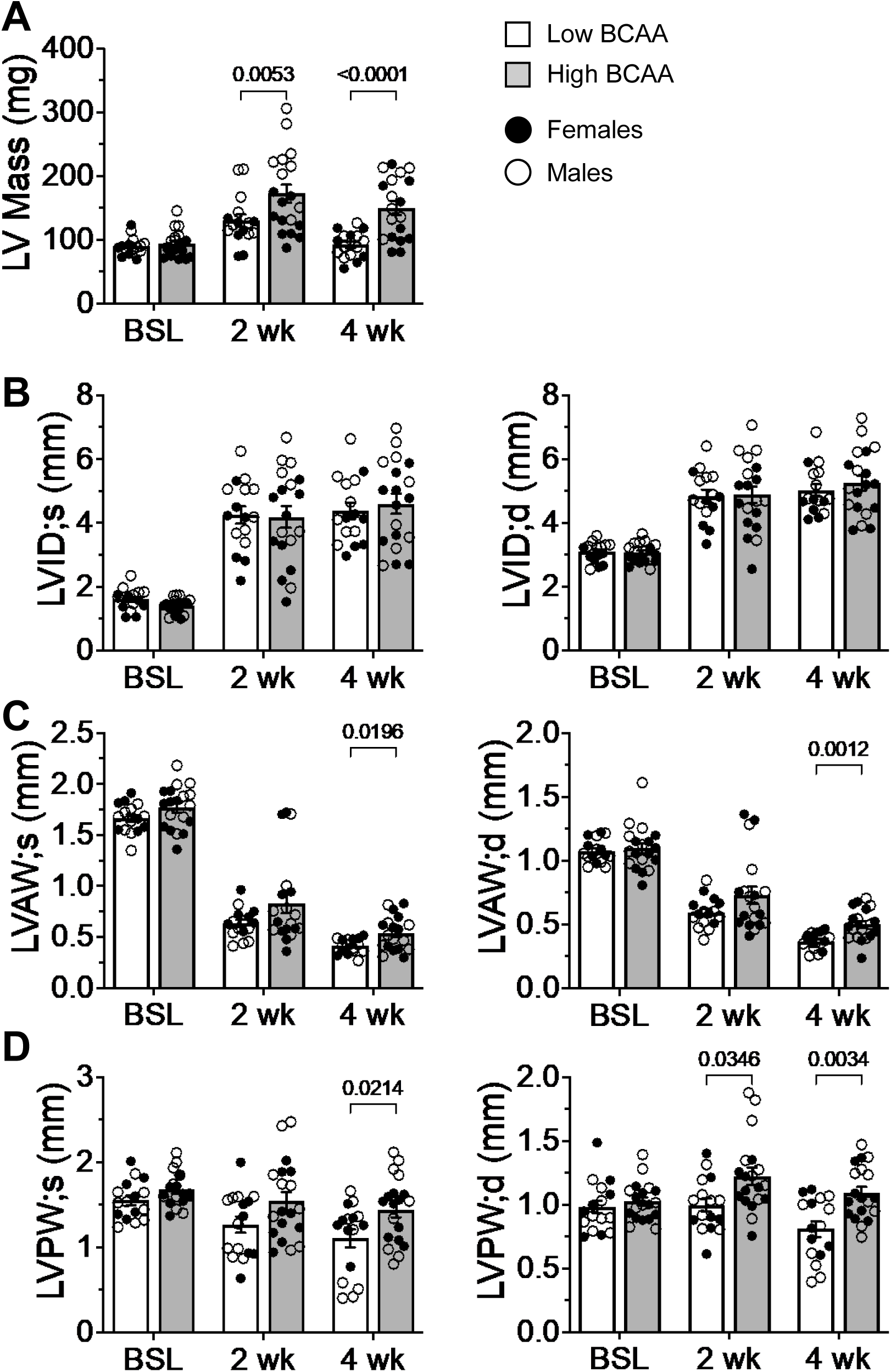
Low dietary BCAA results in smaller LV mass and wall thickness post-MI. Echocardiographic measurements of structural features: (**A**) calculated left ventricular (LV) mass; (**B**) left ventricular internal diameter in systole and diastole; (**C**) left ventricular anterior wall thicknesses in systole and diastole; (**D**) left ventricular posterior wall thicknesses in systole and diastole. n = 16–19/group, males and females pooled. Data are mean ± SEM. Two-way repeated measures ANOVA with Šidák post-hoc.

### The influence of dietary BCAAs on cardiac function and structure post-MI is more prominent in male mice

Given that BCAA levels had more prominent effects on heart size and body mass in male mice (see Fig. 2), we further stratified the echocardiographic analyses based on biological sex. Interestingly, both groups demonstrated lower stroke volume and cardiac output in the high BCAA group compared with the low BCAA group, with more significant changes in the male cohort (**Fig. 5A, 5B**). Because body weights differed in male mice with high BCAA feeding, we also normalized cardiac output values to body weight to obtain cardiac index values. As shown in **Fig. 5C**, mice fed the high BCAA diet showed progressively declining cardiac index values after MI; however, provision of the low BCAA diet completely preserved cardiac index. Stratified analysis of the structural parameters indicates that male mice on the high BCAA diet experienced greater levels of cardiac hypertrophy, as shown by higher LV mass, while this effect was absent in female mice (**Fig. 5D**). Assessment of LV wall thickness further revealed that male mice had significantly larger anterior and posterior wall diameters during diastole, which did not reach significance in female mice (**Fig. 5E, 5F**). These results suggest that biological sex governs responses to dietary BCAAs, with male mice showing a more pronounced response in structural and functional remodeling.

**Figure 5:**
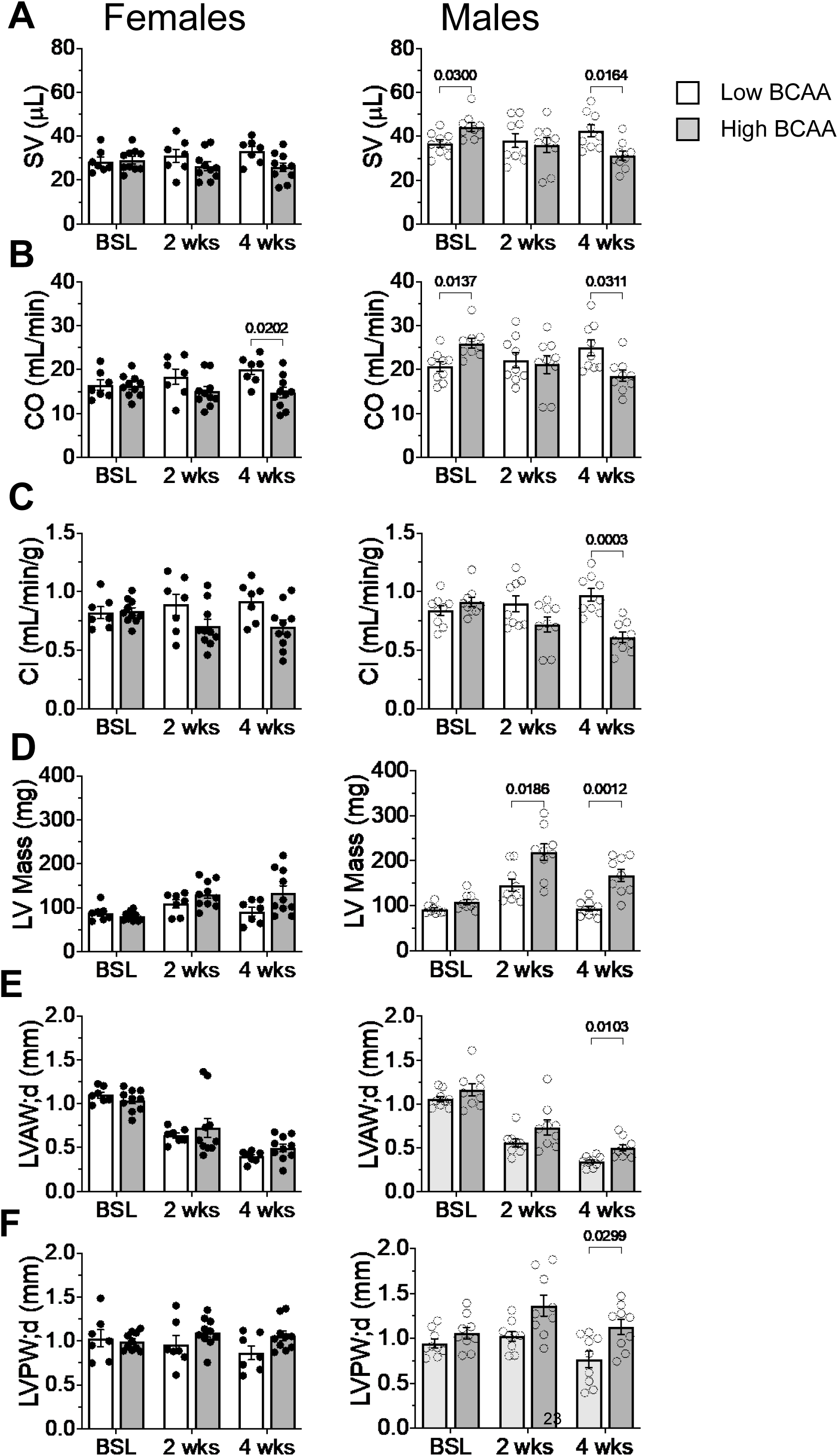
Changes in MI-induced cardiac remodeling with dietary BCAA intervention is more impactful in male mice. Echocardiographic measurements, stratified by sex, for: (**A**) stroke volume (SV); (**B**) cardiac output (CO); (**C**) cardiac index; (**D**) LV mass; (**E**) LV anterior wall thicknesses in diastole; and (**F**) LV posterior wall thicknesses in diastole. n = 7–10/group. Data are mean ± SEM. Two-way repeated measures ANOVA with Šidák post-hoc.

### BCAA levels influence myofibroblast differentiation in vitro

Although dietary BCAAs are known to promote cardiomyocyte hypertrophy,^16^ it is less clear whether they influence fibroblasts in the heart. Therefore, we examined whether BCAAs influence cardiac fibroblast activation *in vitro*. Isolated primary cardiac fibroblasts were treated with either basic fibroblast growth factor (bFGF; control) or transforming growth factor β (TGFβ) in medium containing 0–400 µM BCAAs for 48 h, followed by analysis of Col1a1, periostin, and α-smooth muscle actin (α-SMA) abundance by immunoblotting (**Fig. 6A**). Interestingly, the absence of BCAAs in the culture medium prevented TGFβ-induced upregulation of Col1a1 and periostin (**Fig. 6B, 6C**), but did not prevent upregulation of α-SMA (**Fig. 6D**). Furthermore, intermediate levels of upregulation of Col1a1 and periostin were apparent in fibroblasts cultured in medium containing 100 µM BCAAs, which indicates concentration-dependence in the response. Collectively, these findings indicate that BCAA levels modulate levels of secreted extracellular matrix proteins, but do not affect gross indices of the contractile myofibroblast phenotype, i.e., α-SMA.

**Figure 6:**
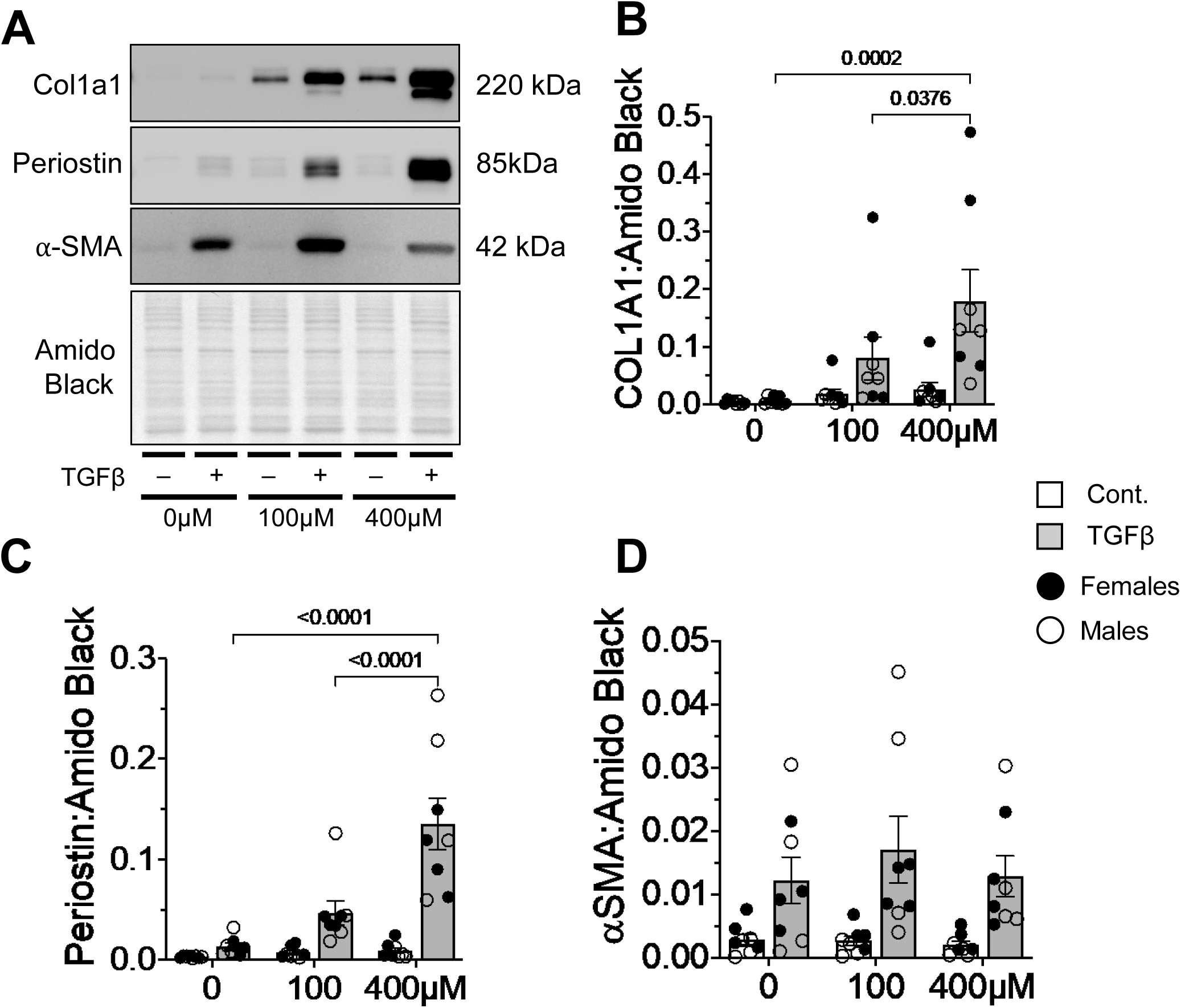
Cardiac fibroblast activation *in vitro* is dependent on BCAA availability. Isolated murine cardiac fibroblasts were treated with TGFβ (or vehicle control) for 48 h in medium containing different concentrations of BCAAs (0 μM, 100 μM, 400 μM) and were assessed for Col1a1, periostin, and α-smooth muscle actin (α-SMA) abundance by immunoblotting. (**A**) Representative image of immunoblots; (**B**) quantification of Col1a1 abundance; (**C**) quantification of periostin abundance; and (**D**) quantification of α-SMA abundance. Proteins were normalized to amido black signal from stained membranes. N = 8/group, with males and females pooled. Data are shown as mean ± SEM. Statistics: two-way ANOVA with Tukey’s post-hoc test.

### High dietary BCAA consumption is associated with eccentric cardiomyocyte hypertrophy and increases in capillary density post-MI in male mice

To determine if circulating BCAA levels influence cardiac fibrosis, we stained cardiac sections with picrosirius red (PSR); however, in contrast to our expectations based on *in vitro* results, we found no difference in total PSR staining between the groups (**Fig. 7A, 7B**). Assessments of transmural scar thickness showed modest differences, with greater thickness in male mice fed the high BCAA diet (**Fig. 7C**). To investigate the potential basis of those observed variations in scar dimensions, infarct regions were imaged via SHG microscopy and subjected to computational enumeration of collagen macrostructural attributes (**Fig. 7D–G**); however, we observed no measurable differences in scar collagen fiber alignment, width, or straightness among the groups (**Fig. 7D–G**). These results suggest that our low BCAA diet may not deplete BCAAs to levels that impede collagen production and scar formation/maturation. Indeed, in a separate experiment, we performed experiments in isolated cardiac fibroblasts using 0, 200, or 400 µM levels of BCAAs, and although complete lack of BCAAs inhibited TGFβ-induced collagen and periostin upregulation, there was no difference between experimental groups that had 200 and 400 µM levels of BCAAs present in the medium. These findings are consistent with the interpretation that the low BCAA diet does not decrease BCAAs below the threshold that impedes extracellular matrix production (**Suppl. Fig. 1**). Further consistent with this interpretation, survival after MI was not different between the low and high BCAA diet groups when either sexes were combined (Kaplan-Meier analysis, p = 0.51, n = 24 per group) or separated by sex (Kaplan-Meier analysis, males p = 0.95, n = 13 per group; females p = 0.26, n = 11 per group), supporting the idea that the low BCAA diet did not affect scar formation or promote cardiac rupture (**Suppl. Fig. 2**).

**Figure 7.**
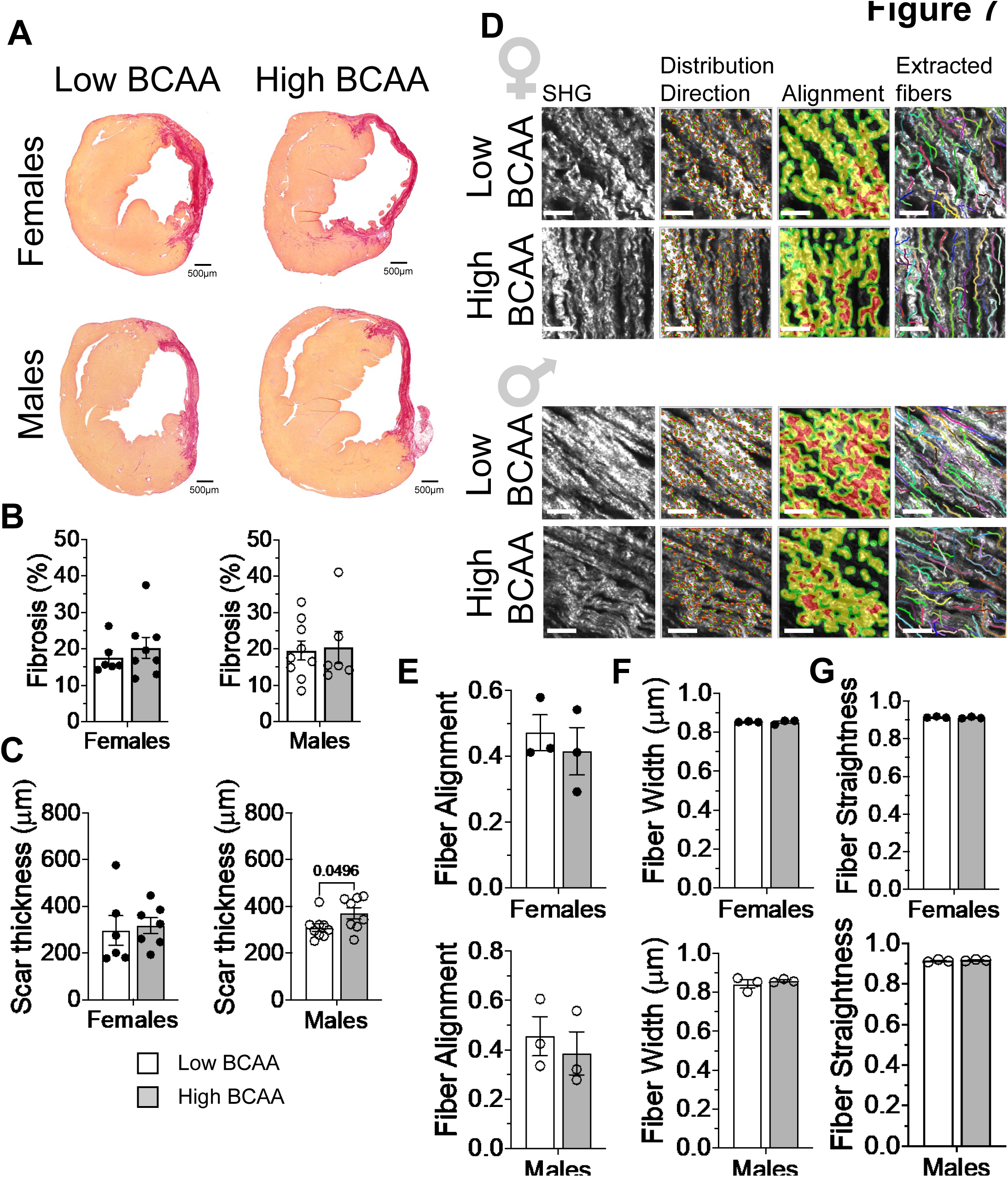
Modified dietary BCAA does not impact cardiac fibrosis and scar remodeling. Examination of cardiac collagen content and organization: (**A**) Intact mid-papillary sections stained with picrosirius red (PSR); (**B**) quantification of PSR signal (cardiac fibrosis) and (**C**) scar width (n = 6–9 biological replicates); (**D**) Computational assessment of scar collagen architecture in second harmonic generation (SHG)-imaged infarct segments. Left to right, representative master SHG images from post infarct scar regions (ROI: 256 × 256 pixel), CurveAlign-generated images displaying the spatial distribution (red dots) and direction (green lines) of collagen fiber segments, corresponding CurveAlign sourced heatmaps depicting the relative alignment of collagen fibers across the region (red denotes regions of high alignment), and, lastly, CT-FIRE extracted fibers overlaid on master SHG images (individual fibers are separated by varying colored lines). From these computational analyses, macrostructural collagen attributes were enumerated and graphed, including, (**E**) fiber alignment (**F**) fiber width, and (**G**) fiber straightness (n = 3 biological replicates per sex per group). Data are mean ± SEM. Normality was assessed via Shapiro-Wilk test. Parametric unpaired student’s t-test or non-parametric Mann-Whitney U test as appropriate. Scale bar = 8.14 μm.

To assess gross differences in cardiomyocytes, we measured cardiomyocyte size and capillary density by staining sections with wheat germ agglutinin (WGA) and isolectin B4. Surprisingly, we found no differences in cardiomyocyte cross-sectional area in hearts from low and high BCAA-fed mice (**Fig. 8A, 8B**); however, WGA assessments of cardiomyocyte length^22, 23^ demonstrated longer myocytes in mice fed the high BCAA diet (**Fig. 8D, 8E**). Of note, capillary density was also observed to be higher in the high BCAA-fed male group, which may suggest a compensatory increase in angiogenesis to support increases in cardiomyocyte volume (**Fig. 8C**). These findings indicate that high dietary BCAA consumption in male mice contributes to eccentric hypertrophy with a compensatory increase in capillary density after MI.

**Figure 8.**
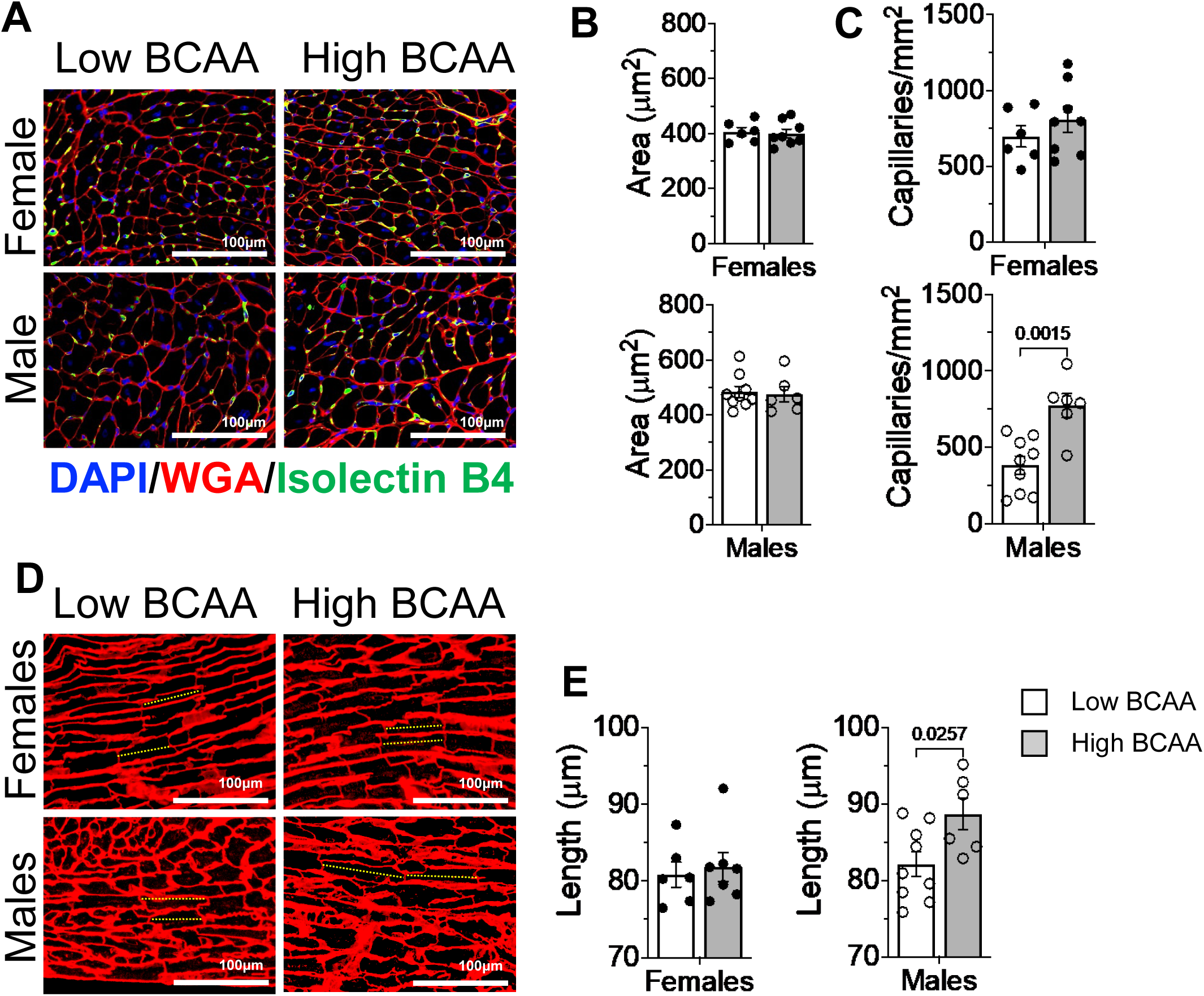
High dietary BCAA consumption is associated with eccentric cardiomyocyte hypertrophy and higher capillary density in male mice. (**A–C**) Cardiomyocyte cross-sectional area (wheat germ agglutin, WGA), and capillary density (Isolectin B4); (**D, E**) Cardiomyocyte length measurements (averaged from 13–73 identified cells per section). Data are mean ± SEM. n = 6–9 biological replicates; Normality was assessed via Shapiro-Wilk test. Parametric unpaired student’s t-test or non-parametric Mann-Whitney U test as appropriate.

## DISCUSSION

Although numerous studies have highlighted a significant role for BCAAs in cardiovascular disease risk,^7, 26^ it remains unclear how levels of circulating BCAAs influence cardiac remodeling after MI. Accordingly, the current study was designed to understand whether modulating BCAA levels via the diet influences MI-induced remodeling. To address this question as simply as possible, we chose diets that are matched for calorific content and nitrogen balance, with the major difference being BCAA levels. Notably, the high BCAA diet and the typical normal chow diet led to similar levels of circulating BCAAs, with the low BCAA diet decreasing BCAA blood levels substantially. Our findings show that provision of a diet low in BCAAs ameliorates cardiac hypertrophy, lowers lung edema, and preserves stroke volume and cardiac output after MI. Moreover, our findings indicate that male mice are more responsive to BCAAs. Collectively, these findings provide support to the concept that simple modifications to the diet could have marked effects on cardiac remodeling after MI and that BCAA levels, in particular, are impactful modifiers of cardiac recovery.

Although diet is a readily modifiable lifestyle factor that could improve cardiovascular outcomes, there remains a paucity of information about what components of the diet may affect cardiac remodeling and function after MI. Early investigations into dietary strategies for post-MI patients revealed a lack of consensus on dietary guidelines,^27^ with initial attempts to optimize dietary intervention leading to broad recommendations such as providing a liquid diet during the first 24–48 h after MI, followed by adherence to low sodium and low cholesterol diets.^28, 29^ More recently, “heart healthy” diets have been reported to be protective against adverse cardiovascular events.^30, 31^ These benefits may partially be attributed to the unique composition of these diets, which are often rich in fruits, vegetables, and legumes.^32, 33^ In particular, Mediterranean dietary patterns have been shown to be modestly protective against MI recurrence and composite cardiovascular disease outcomes.^34^ Studies such as this have led to broad, helpful recommendations that provide general guidance, e.g., implementation of plant-based and Mediterranean-based diets, minimization of saturated and trans fats, higher dietary fiber intake, and diminished consumption of simple carbohydrates.^35, 36^ Nevertheless, it remains unclear which specific components of the diet have deleterious or salutary effects after MI.

Preclinical studies are helpful in shedding light on how particular nutrients, such as BCAAs, affect heart health after cardiac insult or injury. Although the diets used in our study had modified levels of all BCAAs (i.e, leucine, isoleucine, and valine), a previous study in which only dietary leucine was elevated after MI suggests that leucine has generally beneficial effects, attenuating fibrosis and apoptosis and increasing survival.^15^

These findings indicate that individual BCAAs may have different (patho)physiological effects—a claim further supported by recent studies showing that the adverse metabolic effects of BCAAs are mediated by isoleucine and valine, and not leucine.^37^ Nevertheless, provision of a BCAA-free diet to mice with pressure overload significantly attenuated maladaptive fibrosis and cardiac hypertrophy.^17^ Other studies that focused on the effects of BCAAs on the heart show that, in heart failure, cardiac BCAAs are substantially elevated in part due to their impaired catabolism (reviewed in ^6^). For example, direct stimulation of BCAA catabolism with an inhibitor of branched chain ketoacid dehydrogenase kinase decreased cardiac BCAA concentrations and had robust beneficial effects on chronic myocardial remodeling following MI^38^ or pressure overload.^39^ Our findings add to this growing body of literature and show that reducing circulating BCAA concentrations via their dietary restriction has the potential to be a safe and effective therapeutic approach to mitigate pathological cardiac remodeling.

Although the mechanisms by which lowering circulating BCAAs improves remodeling after MI remain unclear, our studies reveal that, compared with the high BCAA diet, the low BCAA diet prevents eccentric cardiomyocyte hypertrophy and improves stroke volume and cardiac output after MI. Previous studies demonstrate that even short-term feeding of the high BCAA diet, in the absence of MI, increases heart size and cardiomyocyte cross-sectional area.^16^ Although we found no differences in cross-sectional area, we show that, after MI, the high BCAA diet significantly increased cardiomyocyte length and increased LV mass in male mice. These findings appear to suggest that a low BCAA may effectively prevent eccentric cardiac remodeling, which could in part underlie observed improvements in stroke volume and cardiac output. Interestingly, both stroke volume and cardiac output were increased at baseline (before MI) in male mice fed the high BCAA diet for 1 week, which suggests that worsened cardiac remodeling after MI in the high BCAA group is the amalgam of two separate hits— elevation of BCAAs and the pathological setting of MI. Although the mechanisms are likely to be multifaceted, they could include BCAA-induced changes in histone propionylation and gene expression,^17^ modulation of mTOR signaling,^40^ cardiac insulin resistance,^7^ and/or extracardiac actions of BCAAs.^38^

Responses to dietary BCAAs are also conspicuously modified by biological sex, with male mice showing more robust responses, which adds another layer of complexity that makes defining a singular mechanism problematic. Notably, only in male mice did high BCAA feeding result in significantly higher body mass and heart weight compared with the low BCAA diet. Given that the diets were matched for calorific content (see Suppl. Table 1) and that previous studies using the same diets and mouse strain found no differences in caloric intake between the two diets,^16^ these findings suggest that BCAAs may provide a higher anabolic stimulus in male mice compared with female mice. Nevertheless, female mice fed the BCAA diets showed trends similar to males, with the low BCAA diet significantly preserving cardiac output and cardiac index. However, remarkably different were the cardiac hypertrophic effects of BCAAs, wherein male mice fed the high BCAA diet and subjected to MI showed higher LV mass, anterior and posterior wall thicknesses, cardiomyocyte elongation, higher capillary density, and worsened lung edema; these effects of high dietary BCAAs were not apparent in female mice. Because increasing BCAA oxidation appears protective and beneficial with respect to cardiac remodeling,^6^ it is tempting to speculate that the sex differences shown here might be due to a heightened capacity of the female heart to oxidize BCAAs, which could be addressed in future studies.

Surprising to us was the finding that the low BCAA diet did not affect myocardial collagen formation or organization *in vivo*. Not only did previous studies using a BCAA-free diet prevent maladaptive fibrosis,^17^ but also our isolated fibroblast studies showed clear evidence that BCAAs are required for collagen and periostin secretion. Interestingly, BCAA availability did not impact TGFβ-induced upregulation of α-smooth muscle actin and acquisition of a contractile phenotype, but it did repress Col1a1 and periostin abundance in a dose-dependent manner, with notable differences discernable when BCAA levels in the medium were decreased to 100 µM. Nevertheless, this effect was lost when BCAA concentrations reached 200 µM (Suppl. Fig 1). Given that the low BCAA diet decreased mean circulating BCAAs to concentrations slightly higher than 100 µM, it is likely that there were sufficient levels of BCAAs to allow for synthesis of collagen. Furthermore, evaluation of collagen fiber macrostructure through SHG imaging within the infarct region revealed no differences in attributes such as collagen fiber alignment, straightness, and width. We hypothesize that there is a threshold level of BCAAs, not reached by the low BCAA diet in this study, at which they may become limiting for synthesis of secreted extracellular matrix proteins.

In summary, we show here that sustained dietary modification of BCAA consumption can alter cardiac remodeling following MI. Specifically, our results indicate that a low BCAA diet mitigates the progressive decline in cardiac output observed with high BCAA diet feeding. This observation may be partially attributed to changes in LV mass and eccentric cardiomyocyte hypertrophy, which is more prominent in male mice. Although the absence of a clear mechanism is a limitation of this study, the discernment of a distinct, singular mechanism is unlikely given the multifaceted actions of BCAAs on anabolic signaling,^40^ post-translational modifications,^17^ insulin sensitivity,^7^ and blood pressure.^38^ Furthermore, it could be conceived that a normal chow group should be included; however, the custom low and high BCAA diets were matched for total protein and nitrogen content as well as carbohydrate and fat content, and the normal chow diet was drastically different, obfuscating comparisons. Moreover, the normal chow diet and the high BCAA diet led to nearly identical levels of circulating BCAAs, which allowed us to address simply the question of whether or not BCAA levels are modifiers of post-MI remodeling. We conclude that dietary BCAAs are relatively strong modifiers of cardiac remodeling post-MI and that a diet low in BCAAs prevents worsening of pathological remodeling after MI. These findings provide an exciting opportunity to further explore how dietary modulation of BCAA levels may be used to improve cardiac function and remodeling after MI.

## Acknowledgments

The authors acknowledge funding support from the National Institutes of Health [HL165813 (D.C.N.), HL168198 (B.G.H.), HL147844 (B.G.H.), HL165826 (C.K.W.), S10OD025178, and P30GM127607] and the American Heart Association [23TPA1141824 (B.G.H.)].

## Author contributions

D.C.N., project planning, execution of experiments, data analysis and interpretation, writing of manuscript, financial support; C.K.W., execution of experiments, data analysis and interpretation, writing of manuscript; M.S.T., execution of experiments, data analysis; Y.M., execution of experiments, data analysis; K.R.B., execution of experiments; R.E.B., execution of experiments; J.B.M., data analysis and interpretation, writing of manuscript; B.G.H., project planning, data analysis and interpretation, writing of manuscript, financial support.

---

**Supplementary Table 1.**

| % by weight | Normal chow | BCAA low | BCAA high |
| --- | --- | --- | --- |
| Protein | 28.7 | 21.5 | 21.5 |
| CHO | 58.2 | 58.6 | 57.2 |
| Fat | 13.1 | 8.1 | 8.1 |
| Kcal/g | 3.0 | 3.9 | 3.9 |

**Supplementary Table 2.**
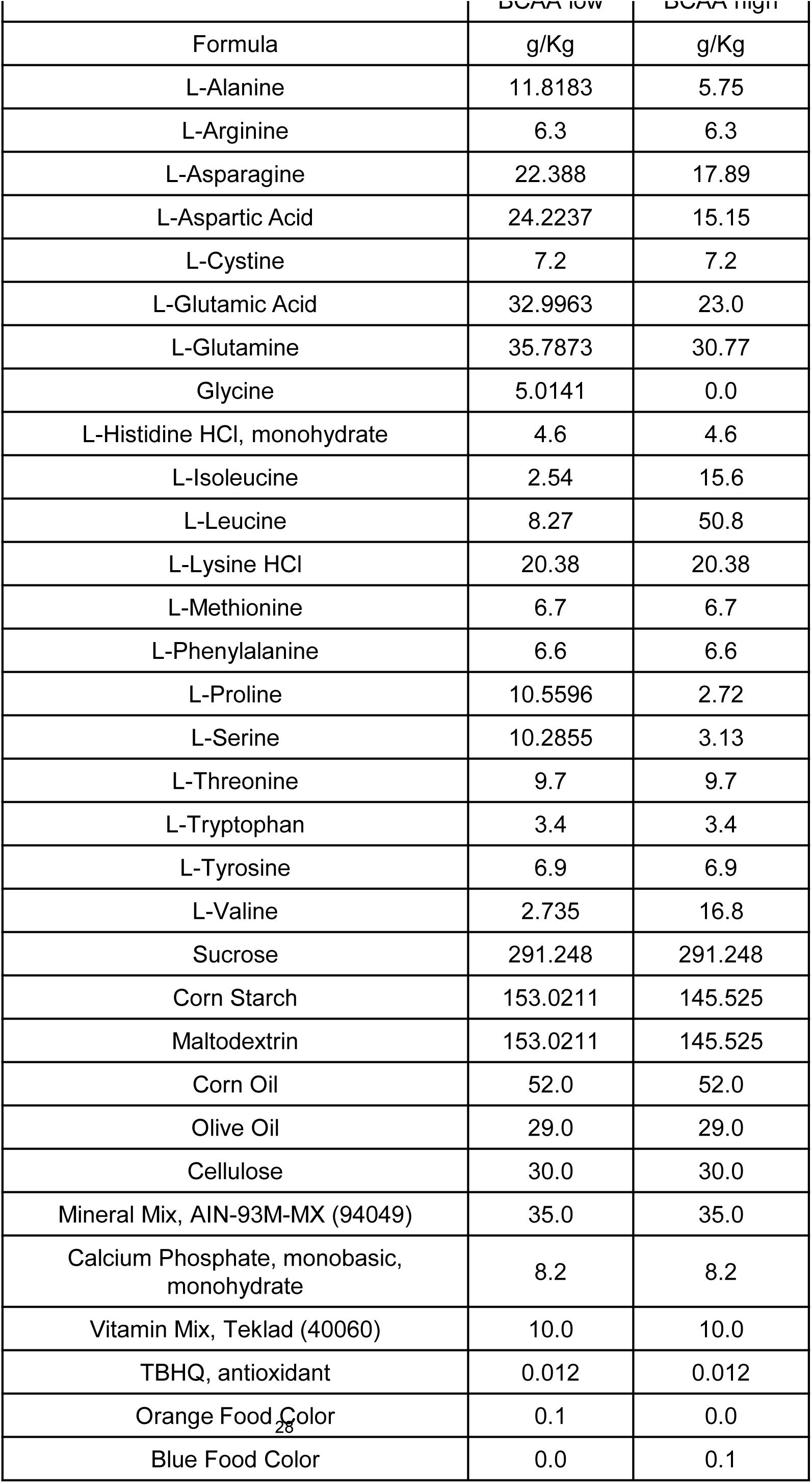

**Supplementary Table 3.**
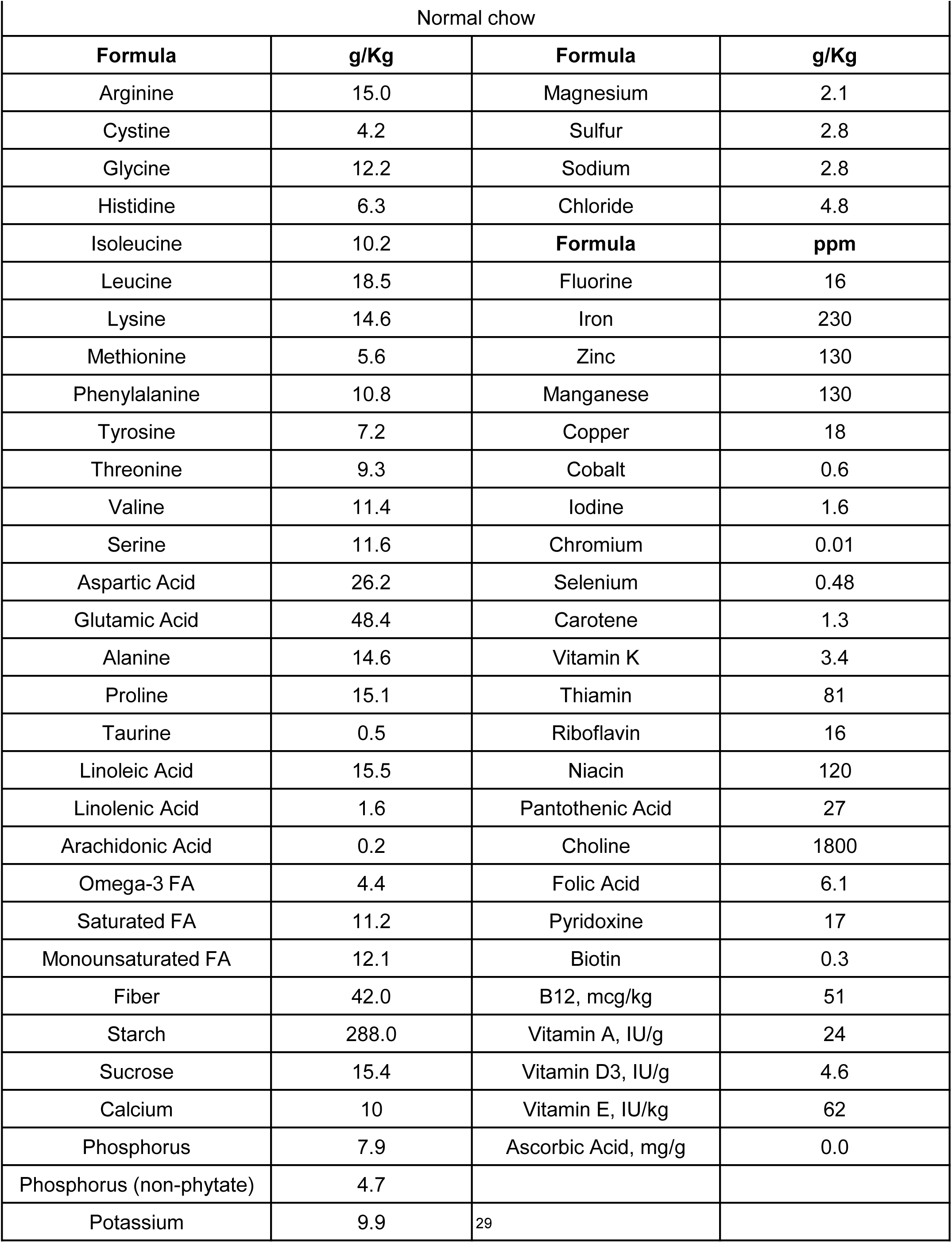

**Supplementary Figure 1.**
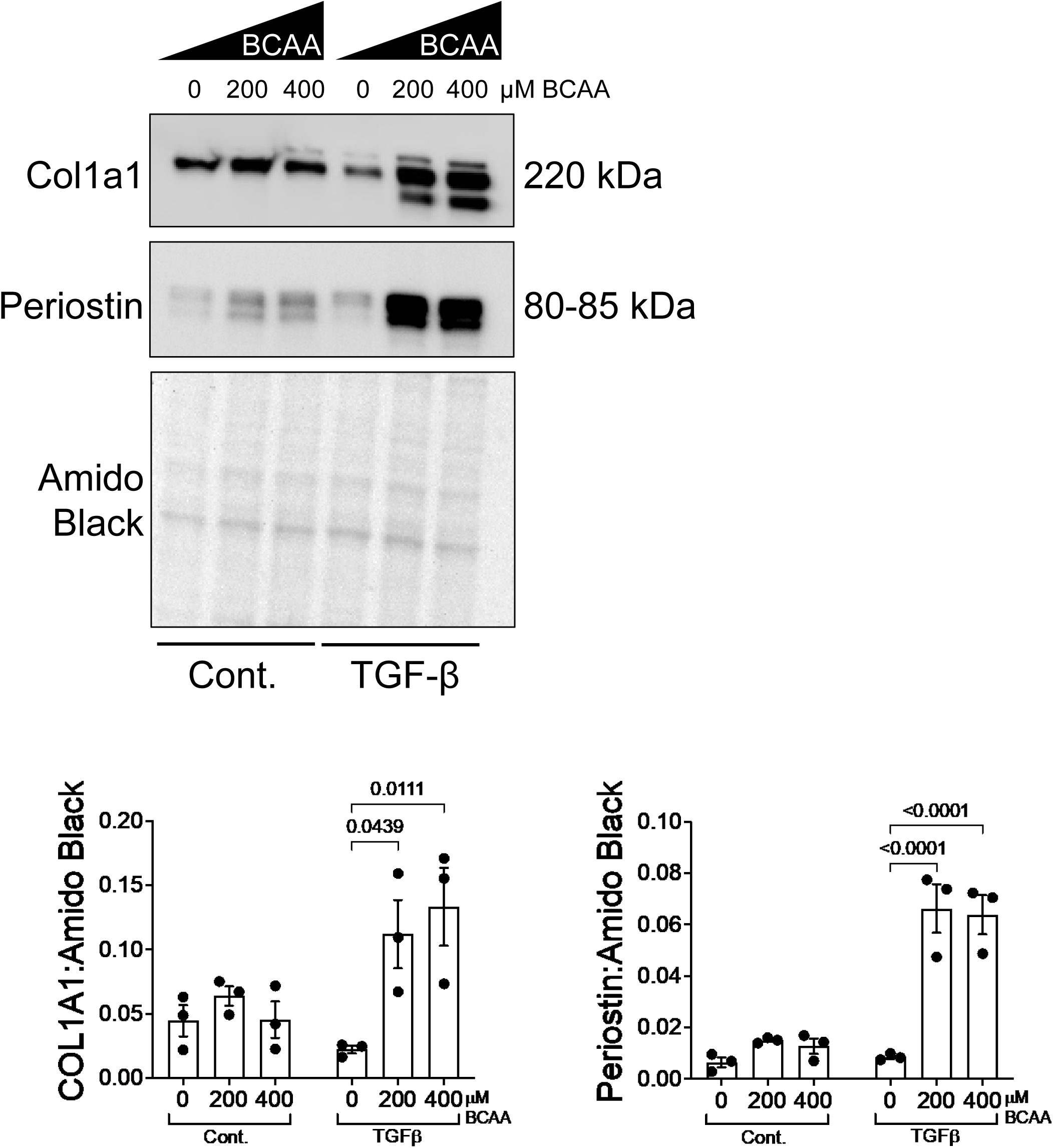
Extracellular BCAA levels of 200 µM or above are sufficient to support TGFβ-induced collagen and periostin secretion. Isolated murine cardiac fibroblasts were treated with TGFβ (or vehicle control) for 48 h in medium containing 0 μM, 200 μM, 400 μM BCAAs. Expression of Col1a1 and periostin was assessed by immunoblotting. Proteins were normalized to amido black signal from stained membranes. n = 3 mice per group. Data are shown as mean ± SEM. Statistics: two-way ANOVA with ^3^T^0^ukey’s post-hoc test.

**Supplementary Figure 2.**
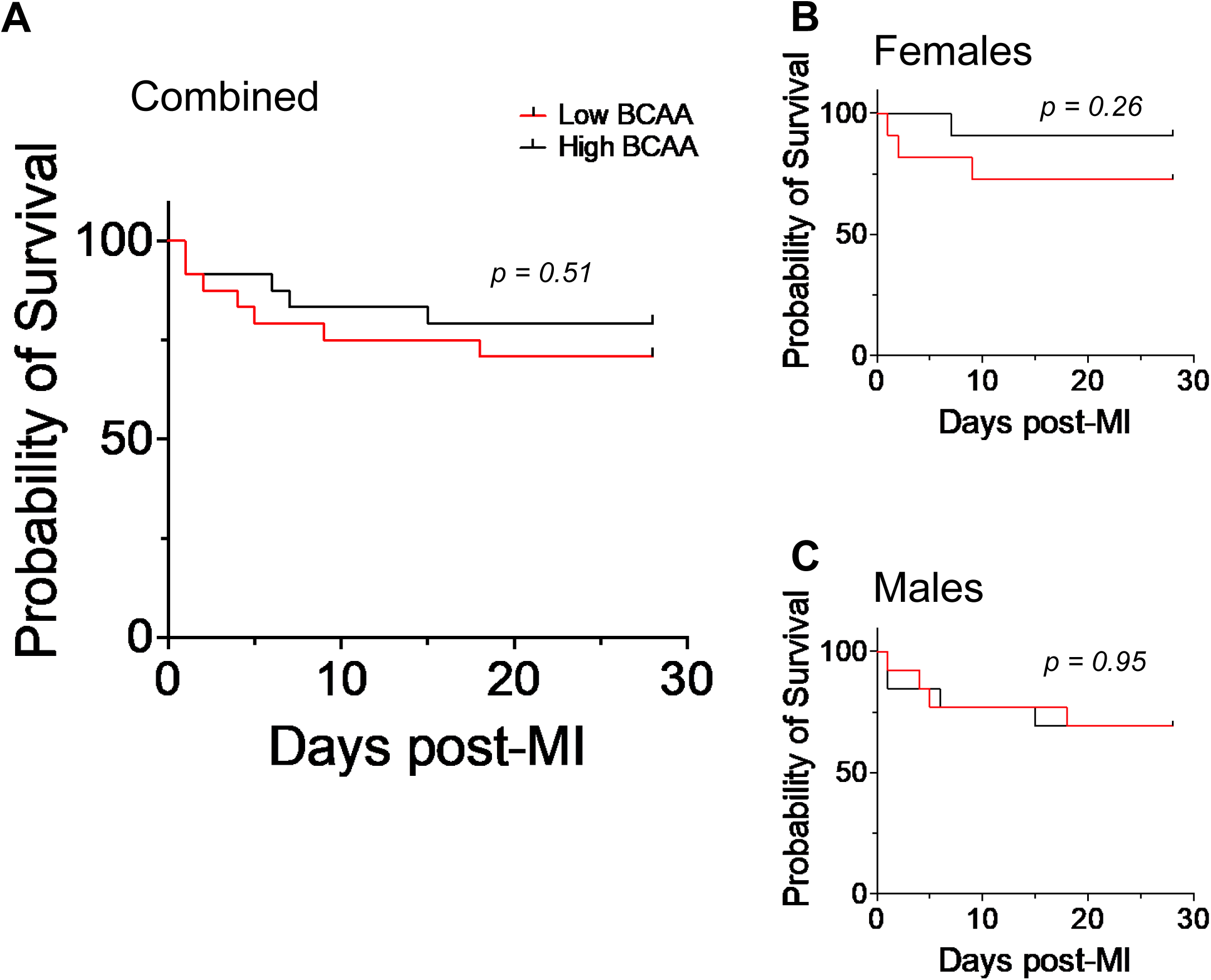
Dietary BCAA intervention does not impact survival post-MI. Kaplan-Meier survival analysis post-MI for combined (A) as well as stratified for females (B) and males (C). n = 11 – 13 per group/sex

